# Neural Responses to Heartbeats of Physically Trained and Sedentary Young Adults

**DOI:** 10.1101/156802

**Authors:** Pandelis Perakakis, Antonio Luque Casado, Luis Ciria, Plamen Ivanov, Daniel Sanabria

## Abstract

Regular physical exercise has a positive impact on brain function and cognitive performance. However, it is not yet clear whether the physiological and behavioral benefits associated to physical exercise are caused exclusively by changes in cardiovascular fitness. Here, we explore the relation between regular physical exercise and transient electroencephalographic responses to afferent cardiac signals. We find differences in the neural processing of heartbeats between individuals who exercise regularly and their sedentary counterparts. These differences, localised at two distinct spatio-temporal clusters, occur before the presentation of a target stimulus and correlate with behavioral performance only in the high-fit group. We hypothesise that the different neural processing of afferent cardiac activity by physically trained individuals reflects enhanced interoceptive sensitivity, which contributes to improved sustained attention. Our results are in line with recent research highlighting the role of neural monitoring of visceral signals in perceptual processing and even the generation of the sense of self.

## Introduction

In the past decades we have learned a lot about the physically active brain. Findings from human and animal studies have shown that regular physical exercise reduces the risk of diseases associated with cognitive deterioration, protects the brain from the adverse effects of ageing, and helps improve cognitive performance (Voss et al., 2013; Hillman et al., 2008). An important question that remains open is whether the brain and cognitive improvements attributed to physical exercise are exclusively mediated by gains in cardiovascular fitness. Although several studies have found a dose-dependent relationship between increases in fitness and brain outcomes (e.g., Erickson et al. 2011; Voss et al. 2012), other studies (e.g., Barrientos et al. 2011; Voss et al. 2010), including two meta-analytic reviews (Etnier et al. 2006; Angevaren et al. 2008), failed to detect a correlation between changes in cardiovascular fitness and cognitive improvements. This uncertainty around the neurophysiological mechanisms underlying the exercise-brain-cognition relationship makes it difficult to evaluate existing and design new cognitive training programs, and to inform public health recommendations. In this study, we investigate the differences in neural monitoring of afferent cardiac information between physically trained and sedentary young adults, and formulate the novel hypothesis that exercise may improve human brain and cognitive function by augmenting interoceptive sensitivity (IS).

Accumulating evidence indicates that the capacity to sense the physiological condition of the body, defined as “interoception”, plays a crucial role in cognitive function and associated behavioural choices. Research in this area, focusing mostly on cardiac perception, has shown that individual sensitivity to interoceptive stimuli modulates implicit memory (Werner et al., 2010), selective and divided attention (Matthias et al., 2009), intuitive decision making (Dunn et al., 2010), visual awareness (Park et al., 2014; Salomon et al., 2016), memory encoding (Garfinkel et al., 2013), and somatosensory processing (Edwards et al., 2009; Gray et al., 2009). Furthermore, neuroimaging studies revealed that the neural systems supporting interoceptive awareness, namely the anterior insular cortex (AIC) and the anterior cingulate cortex (ACC) (Critchley et al., 2004; Gray et al., 2009), are also involved in selective attention (Weissman et al., 2006), stimulus-independent thought (Mason et al., 2007), perceptual decision making (Ploran et al., 2007; Thielscher and Pessoa, 2007), recognition memory (Yonelinas et al., 2005) and executive cognitive control (Cole and Schneider, 2007; Dosenbach et al., 2007; Brass and Haggard, 2007; Eichele et al., 2008; Sridharan et al., 2008).

Research on cardiac interoception further revealed that people vary significantly in their ability to perceive their own heartbeats (Critchley et al., 2004; Herbert et al., 2007; Pollatos et al., 2007; Schandry, 1981). Individual differences in IS are usually assessed objectively by heartbeat detection tasks but are also reflected in the amplitude of the heartbeat-evoked potential (HEP), which quantifies the unconscious neural processing of afferent cardiac signals and is obtained by averaging electroencephalographic activity time-locked to individual heartbeats (Pollatos and Schandry, 2004). Critically, cardiac interoception is not a stable individual trait but can be increased with biofeedback training (e.g., Katkin et al. 1981; Grigg and Ashton 1982; Davis et al. 1986), while training-induced increases in IS are also captured by changes in the amplitude of the HEP (Schandry and Weitkunat, 1990; Canales-Johnson et al., 2015).

The finding that IS can be potentiated through feedback, signals physical exercise as a possible interoceptive training mechanism. Indeed, early research exploring the effect of exercise on interoception reported that heartbeat perception improves during or immediately after moderate or intense physical exercise (Borg and Linderholm, 1967; Montgomery and Jones, 1984; Yuan et al., 2007; Rädler and Schandry, 1988), probably as a consequence of increases in stroke volume and/or heart contractility (Schandry et al., 1993). At the same time, individuals with high aerobic fitness demonstrate better heartbeat perception than those with less or no training (Jones and Hollandsworth, 1981; Schandry et al., 1993). In addition, more recent neuroimaging research has shown that physical exercise produces alterations in the ACC, which is, as mentioned above, a crucial brain structure for interoceptive awareness, and is also related to improved cognitive performance, especially in tasks requiring executive control (Hillman et al., 2008; Colcombe et al., 2004; Chaddock et al., 2012).

Taken together, these findings seem to corroborate the hypothesis that physical exercise may be improving cognitive function and performance by increasing IS. To explore this novel hypothesis, we tested differences in the neural processing of afferent cardiac signals between a group of trained athletes and a group of sedentary individuals, during the performance of a 60 — min Psychomotor Vigilance Task (PVT). In previous work with the same participants we investigated the behavior of separate brain (event-related potentials, time-frequency responses) and cardiac (transient heart rate deceleration) physiological parameters (Luque-Casado et al., 2016; Casado et al., 2016). Here, we turn to a measure of brain-heart interaction that has never been investigated in this context. Specifically, we look at differences in the HEP amplitude evoked by the first two heartbeats following the warning signal of the PVT task. Further, to explore the relationship between IS and cognitive performance, we look at the correlation between HEP amplitude and reaction times to the subsequent target stimulus.

## Materials and Methods

### Participants

Fifty young male adults with normal or corrected-to-normal vision and no clinical history of cardiovascular or neuropsychological disorders participated in the study. Twenty-five of the participants, who were assigned to the “high-fit” group, were recruited from local triathlon clubs (N = 15) and from the Faculty of Physical

Activity and Sport Sciences of the University of Granada (N = 10). Another twenty-five undergraduate University students with sedentary lifestyle were assigned to the “low-fit” group. The participants in the high and low-fit groups met the inclusion criteria of reporting at least *8h* of training per week or less than 2h, respectively. The size of the sample was determined by an a priori power analysis indicating a minimum of 22 participants per group for a power level of 0.80. The experiment was conducted according to the ethical requirements of the local committee and in compliance with the Helsinki Declaration. All participants signed an informed consent and were paid for their participation. They were required to maintain a regular sleep-wake cycle for at least one day before the study and to avoid caffeine and vigorous physical activity before the visit to the laboratory. All data presented here were analysed and reported anonymously. Five participants from the high-fit group and eight from the low-fit group were excluded from the analysis due to noisy signals (see *Electrophysiological recordings and analysis).*Table 1 presents the anthropometrical characteristics of the 37 participants (20 high-fit and 17 low-fit) whose data were analysed.

**Table 1:** Mean and 95% Confidence Interval (CI) of descriptive, fitness, behavioral and EEG data for the high-fit and low-fit groups. Only data of the participants included in the analyses are reported. BMI = Body Mass Index; VO_2_ = Oxygen Consumption; VAT = Ventilatory Anaerobic Threshold; RT = Reaction Times; HEP = Heartbeat-Evoked Potential

|  | High-fit | Low-fit |
| --- | --- | --- |
| Anthropometrical characteristics |  |  |
| Sample size | 20 | 17 |
| Age (years) | 22.55 ± 1.7 | 22.94 ± 0.9 |
| Height (cm) | 1.77 ± 0.02 | 1.78 ± 0.03 |
| Weight (kg) | 69.28 ± 2.82 | 75.18 ± 8.7 |
| BMI($kg/m^2$) | 22.15 ± 0.8 | 23.66 ± 2.25 |
| Fitness test parameters |  |  |
| VO<sub>2</sub> at VAT ($mL/(kg \times min)$) | 44.59 ± 3.6 | 19.48 ± 2.6 |
| Relative power output at VAT ($W/kg$) | 3.55 ± 0.3 | 1.39 ± 0.2 |
| Behavioral & EEG responses |  |  |
| RT Block 1 (ms) | 267.98 ± 9.14 | 282.8 ± 16.7 |
| RT Block 2 (ms) | 291.05 ± 12.6 | 297.52 ± 19.38 |
| HEP amplitude – positive cluster ($\mu V$) | 1.2 ± 0.26 | 0.06 ± 0.27 |
| HEP amplitude – negative cluster ($\mu V$) | -0.67 ± 0.58 | 0.1 ± 0.55 |

### Psychomotor vigilance task

We used a modified version of the Psychomotor Vigilance Task (PVT; Wilkinson and Houghton 1982) to measure sustained attention by recording participants’ reaction time (RT) to visual stimuli that occurred at random inter-stimulus intervals (Basner and Dinges, 2011). Each trial began with a blank screen in a black background for 2s, followed by the presentation of an empty red circumference (6.68° x 7.82°). After a random time interval (between 2 and 10s), the circumference was filled in a red color. Participants were instructed to respond as fast as possible after the circle was colored. The colored circle was displayed for 0.5s and participants had a maximum of 1.5s to respond. They had to respond with their dominant hand by pressing the space bar on a PC keyboard. A visual feedback indicating their RT was displayed for 0.3s after the response, except in the case of an anticipated response (“wait for the target”) or if no response was made within 1s after target offset (“you did not answer”). The next trial began immediately after feedback offset (see Fig. 1a). Response anticipations were considered as errors. The task comprised a single block of *60min* and the mean number of trials per participant was 402 ± 8.9.

**Figure 1:**
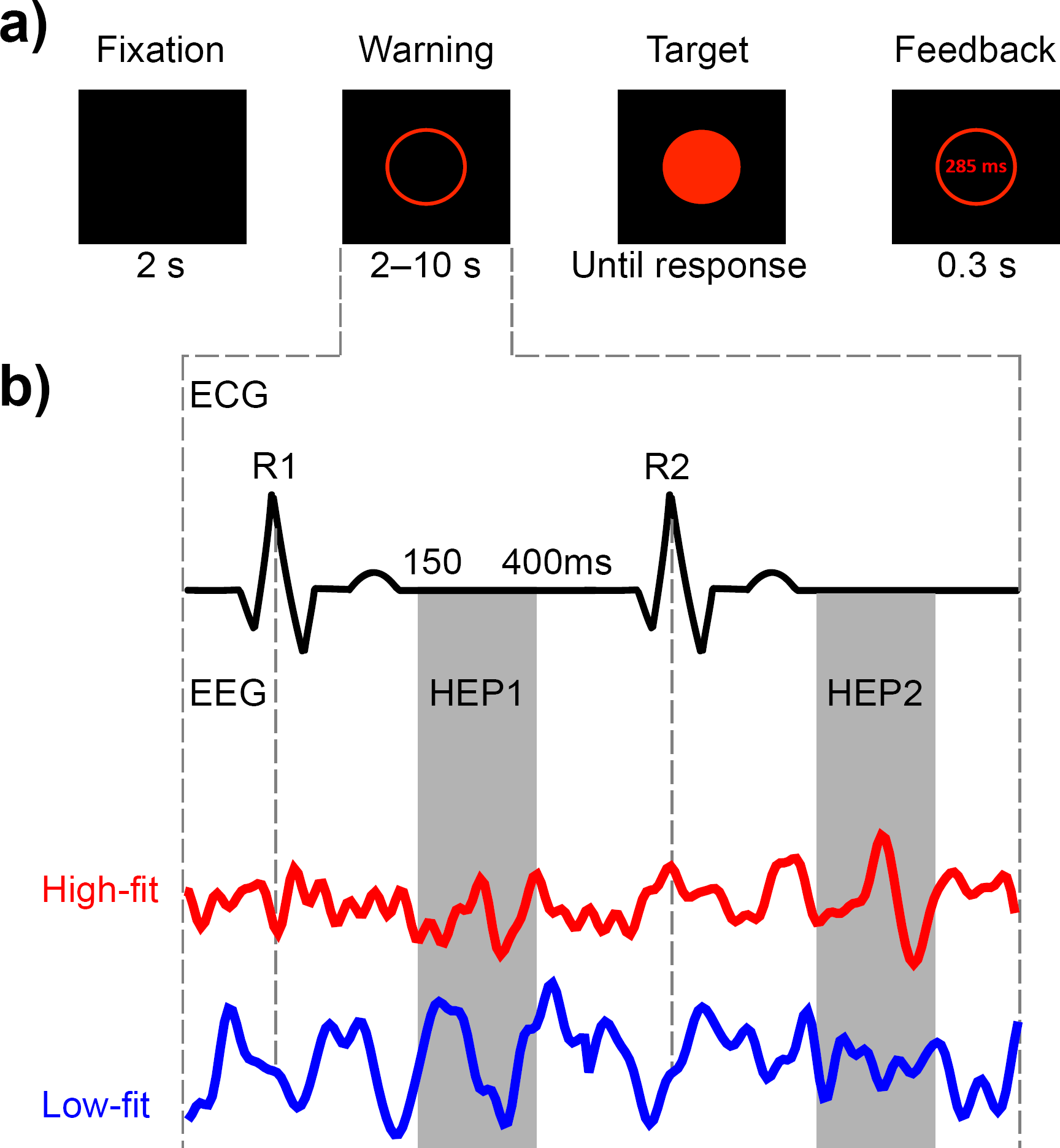
Experimental design and rationale. (**a**) Time course of a single PVT trial. Participants had to respond as soon as possible after the appearance of the Target stimulus. (**b**) The first two cardiac cycles occurring after the Warning stimulus were used for computing the HEP. The EEG signal was averaged locked to the two post-warning R-peaks (R1, R2), resulting in two distinct event-related components (HEP1 and HEP2). The statistical analysis to test differences between the High-fit and Low-fit groups focused on the time window during cardiac relaxation that is free from cardiac artifacts (approximately 150-400ms after the R-peak).

### Incremental effort test

A brief anthropometric study of each participant was performed to measure height, weight, and body mass index (see Table 1). We used a ViaSprint 150 P cycle ergometer (Ergoline GmbH, Germany) to induce physical effort and to obtain power values, and a JAEGER Master Screen gas analyzer (CareFusion GmbH, Germany) to provide a measure of gas exchange during the effort test. Before the start of the test, participants were fitted with a Polar RS800 CX monitor (Polar Electro Oy, Kempele,Finland) to record their HR during the incremental exercise test and the cycle ergometer was set to the individual anthropometric characteristics (Luque-Casado et al., 2013). The incremental effort test started with a 3 — min warm-up at 30W, with the power output increasing 10W every minute. The test began at 60W and was followed by an incremental protocol with the power load increasing 30W every *3min.* Workload increased progressively during the third minute of each step *(5W* every 10s). Therefore, each step of the incremental protocol consisted of *2min* of stabilized load and 1min of progressive load increase. Each participant set his preferred cadence (60 — *90rpm)* during the warm-up and was asked to maintain this cadence throughout the protocol. The ergometer software was programmed to increase the load automatically. Determination of the ventilatory anaerobic threshold (VAT) was based on the respiratory exchange ratio (RER) [RER = CO_2_ production / O_2_ consumption]. More specifically, VAT was defined as the VO_2_ at the time when RER exceeded the cutoff value of 1.0 (Davis et al., 1976; Yeh et al., 1983). The researcher knew that the participant had reached his VAT when the RER was equal to 1.00 and did not drop below that level during the 2 — *min* constant load period or during the next load step, never reaching the 1.1 RER. The submaximal incremental test ended once the VAT was reached. Oxygen uptake *(mL/(min × kg)),* RER, relative load *(W/kg),* and HR (bpm) were continuously recorded during the entire incremental test. This dataset determined the fitness level of each participants (see Table 1).

### Experimental procedure

Upon arrival to the laboratory, participants were seated in front of a 19 — *inch* PC monitor in a dimly-illuminated, sound-attenuated room with a Faraday cage. The centre of the PC screen was situated at eye level approximately 60*cm* from the head of the participant. A PC keyboard was used to record behavioral responses. After a brief training session participants were instructed to complete a 60 — min version of the PVT (Fig. 1a). Upon task completion, all participants performed the incremental effort test to objectively evaluate their fitness level (Table 1). The experimental session was administered during daylight hours, with approximately half the participants coming to the lab in the morning and the other half in the afternoon.

### Electrophysiological recordings and analysis

Continuous EEG signals were recorded at 1024H*z* using a 64-channel BioSemi Active Two amplifier system (Biosemi, Amsterdam, Netherlands). ECG signals were simultaneously recorded using two FLAT active electrodes (Ag/AgCl; Biosemi, Amsterdam, Netherlands) arranged at a modified lead I configuration (i.e., right and left wrists). The R-peaks of the QRS ECG complex were first automatically detected by the ECGLAB matlab software (De Carvalho et al., 2002) and then visually inspected for manual artifact correction. A total of 6 participants (2 high-fit and 4 low-fit) were excluded from posterior analysis due to a noisy ECG signal. EEG preprocessing was conducted using custom matlab scripts and the EEGLAB (Delorme and Makeig, 2004) and Fieldtrip (Oostenveld et al., 2011) Matlab toolboxes. EEG data were resampled at *256Hz* and bandpass filtered offline from 0.5 to 40H*z*. An independent component analysis was used to identify and remove EEG components reflecting eye blinks (Delorme and Makeig, 2004).

For HEP analysis, the EEG signal was segmented to 800 — ms epochs extending from —200ms to 600ms time-locked to the first and second R-peak following the warning stimulus. We focused only on the first two cardiac cycles following the warning signal to avoid an overlap with the target stimulus (see Fig. 1b). EEG epochs were baseline-corrected relative to a time window extending from —200ms to —50ms, to exclude the rising edge of the R-peak (Canales-Johnson et al., 2015). Trials that contained fluctuations exceeding ±200μV were rejected. To ensure an adequate signal-to-noise ratio and to reduce type I error inflation by post-hoc exclusion of subjects, we set an a priori criterion of excluding participants for whom more than 25% of trials were rejected (Luck, 2014). Based on this criterion, another 7 participants (3 high-fit and 4 low-fit) were excluded from further analysis.

### Statistical analysis

The behavioral data were analyzed using a repeated measures analysis of variance (ANOVA) with the between-participants factor of Group (High-fit, Low-fit) and the within-participants factor of Block (B1, B2). We split the 1 — hr PVT to two 30 — min blocks to investigate the effect of task duration (time-on-task), reported in our previous studies with the same participants (Luque-Casado et al., 2016; Casado et al., 2016). Effect sizes are reported by partial eta-squared *(npartial*^2^). A t-test for independent samples was applied to analyze the participants’ fitness data.

We tested the significance of HEP differences between the high-fit and low-fit groups using a cluster-based non-parametric permutation test (Maris and Oostenveld, 2007) as implemented in the Fieldtrip toolbox. This test does not require the definition of a priori spatial or temporal regions of interest and corrects for multiple comparisons in space and time. For each electrode × time pair a t-test for independent samples was performed and pairs with positive and negative t-values that exceeded a threshold *(p* < 0.05, two-tailed) were clustered together based on temporal and spatial adjacency. Cluster-level statistics were then calculated by taking the sum of the *t*-values within each cluster. The trials from the two datasets were then randomly shuffled and the maximum cluster-level statistic for these new shuffled datasets was calculated. The above procedure was repeated 4,000 times to estimate the distribution of maximal cluster-level statistics obtained by chance. The proportion of random partitions that resulted in a larger test statistic than the original one, determined the two-tailed Monte-Carlo *p*-value.

## Results

### Descriptive and fitness data

The t-tests for independent samples revealed significant differences between groups in the VO_2_ at VAT, t(35) = 10.74, *p* < 0.01, and the relative power output at VAT, t(35) = 11.09, *p* < 0.01. All anthropometric data indicated a marked difference in the fitness level between groups (see Table 1).

### Behavioral results

A repeated-measures ANOVA with the between-participants factor of Group (High-fit, Low-fit) and the within-participants factor of Block (B1, B2) was conducted on participants’ mean RTs. The main effect of Block reached statistical significance, 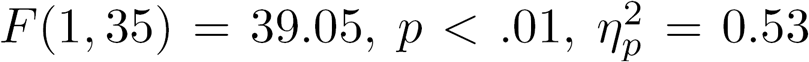, but neither the main effect of Group nor the interaction between Block and Group showed significant differences (both ps ≥ 0.18). This result shows a similar tendency but does not precisely replicate the interaction effect we reported in previous studies with the same participants (Luque-Casado et al., 2016; Casado et al., 2016). Two issues may account for this discrepancy: 1) we used only two 30 — *min* blocks instead of five 12 — min blocks, and 2) due to EEG artifacts related to the novel analysis reported here, not exactly the same participants were analysed.

### HEP results

We computed the HEP time-locked to the first and the second R-peaks following each warning stimulus appearing prior to the target stimulus of a psychomotor vigilance task taxing sustained attention. We then averaged HEPs from the high-fit (HF) experimental group during the first (B1) and the second (B2) 30 — *min* blocks of the task. HEPs from the first (HEP1) and the second (HEP2) cardiac cycles after the warning stimulus were averaged separately (see Fig. 1b). The same procedure was repeated for the low-fit (LF) group, resulting in a total of 8 datasets: 2 Groups *(HF, LF*) × 2 post-warning cardiac cycles (*HEP* 1, *HEP2*) × 2 temporal blocks (B1,B2).

We first tested the interaction Group x Block by submitting the 150 — 400ms post R-peak time window of the HEP difference *B1 — B2* to the cluster-based permutation test (see Statistical analysis section). This time window was chosen based on the latency of HEP differences in a previous study with similar methodology where the effect of feedback training on HEP was tested (Canales-Johnson et al., 2015), and also because it corresponds to the heart relaxation period where EEG responses to heartbeats are free from electrical artifacts caused by heart contractions (Gray et al., 2007). The test did not yield any significant clusters neither for HEP1 nor HEP2. We then turned to the main effect of Group. HEP amplitude differed significantly between the high-fit and low-fit groups over frontocentral (Monte-Carlo p = 0.02) between 257 and 358*ms* post R-peak (Fig.2a), and left occipitoparietal (Monte-Carlo *p* = 0.008) between 253 and 343*ms* post R-peak (Fig. 2b). These clusters were present only in HEP2, the HEP computed for the second cardiac cycle after the warning stimulus. In Figure 2, we observe that HEP values were lower for the high-fit group in the frontocentral cluster and higher in the the occipitoparietal cluster.

**Figure 2:**
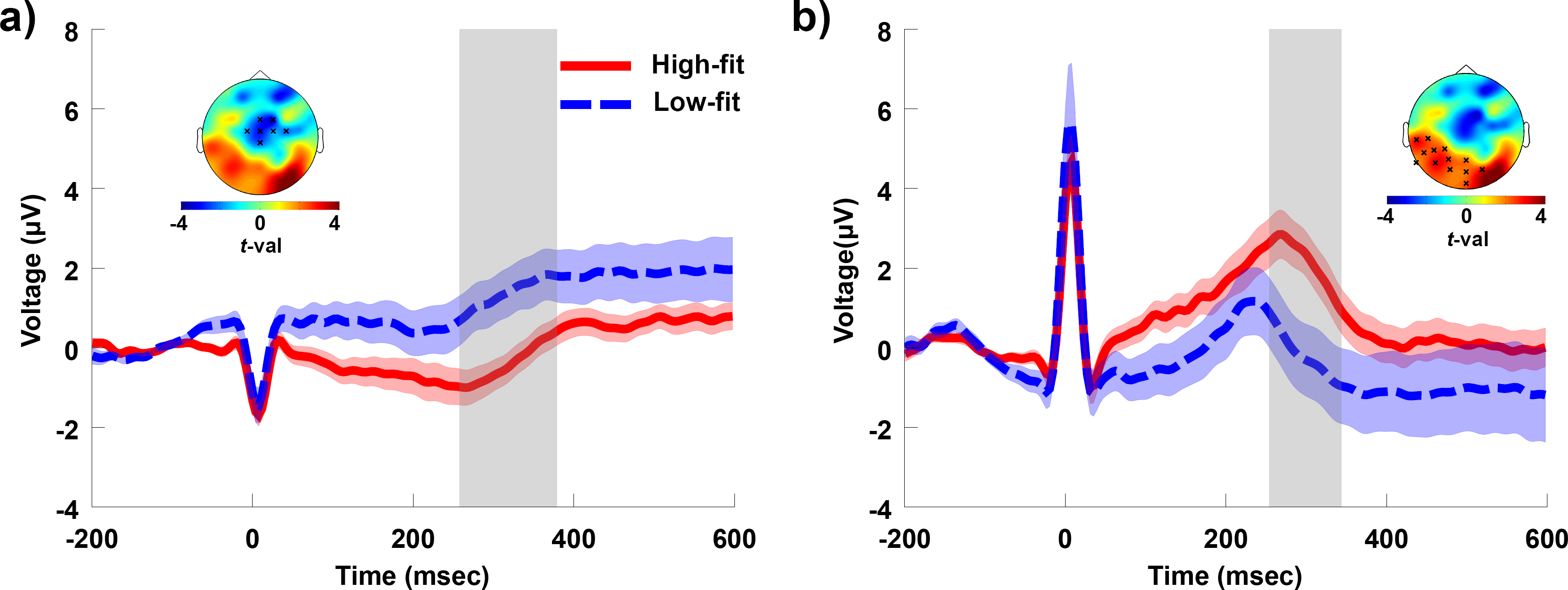
EEG responses locked to the second heartbeat after the Warning stimulus (HEP2). (**a**) Grand average results for the frontocentral cluster indicated by the black “x” marks at the topographical map. Significant differences (Monte-Carlo p=0.02) were identified across the cluster in the time window indicated by the shaded area (between 257 and 358 ms). (**b**) Grand average results for the occipitoparietal cluster indicated by the black “x” marks at the topographical map. Significant differences (Monte-Carlo p=0.008) were identified across the cluster in the time window indicated by the shaded area (between 253 and 343 ms).

Next, we investigated whether the amplitude of neural responses to heartbeats is associated to the reaction times to the target stimulus. In each subject, we sorted single trials according to the mean amplitude of the HEP computed at the electrodes of each cluster and in their corresponding significant time windows. We then split sorted trials into three bins of low, medium and high HEP amplitude. We calculated for each bin the change in the reaction time relative to the subject’s mean reaction time across all trials, and computed the linear regression between HEP amplitude and reaction time change (for an example of a similar procedure see Park et al. 2014). We performed this analysis separately for each significant cluster and for each one of the groups. Across high-fit subjects, slope values of the linear regressions in the occipitoparietal cluster were significantly larger than 0 (mean slope,-0.26 ± 0.52, t =-2.19, p = 0.04). Pearson correlations showed a similar, but less significant trend (mean Pearson r,-0.04 ± 0.09, t =-1.84, p = 0.08; Fig. 3a). For the low-fit group, neither slope values nor Pearson correlations were significantly larger than zero (mean slope, 0.08 ± 0.38, t = 0.82, p = 0.43; mean Pearson r, 0.005 ± 0.38, t = 0.47, p = 0.64; Fig. 3b). No significant deviations from zero were found for the slopes or for Perason correlations at the frontocentral cluster, either for the high-fit or the low-fit group.

**Figure 3:**
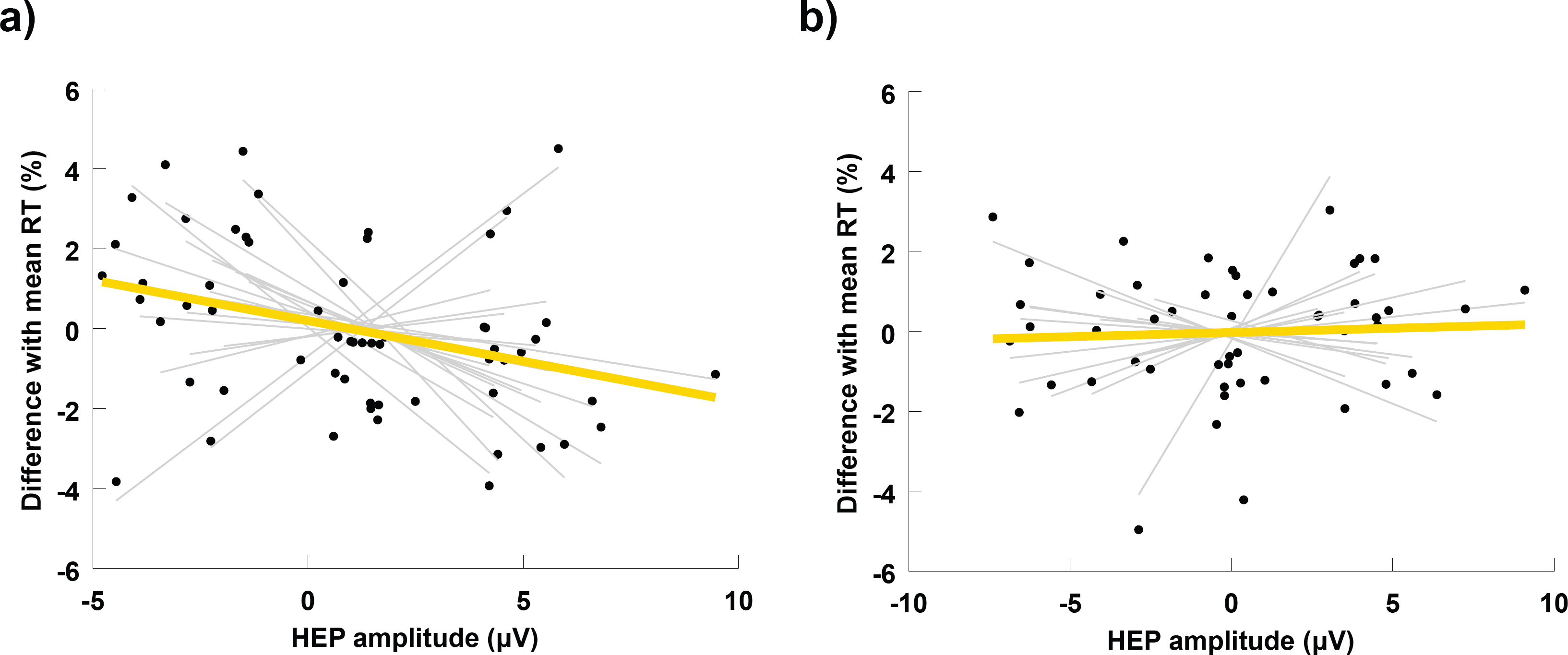
Correlations between heartbeat-evoked potentials and reaction times. In each subject, trials were sorted in three bins according to the amplitude of the EEG response to heartbeats. The position of each black dot in the horizontal axis represents the mean HEP amplitude for a given bin, and its position in the vertical axis the mean reaction time for the same bin, expressed as a percentage difference with the mean reaction time across all trials of the same subject. Thin grey lines represent the slope of the linear regression between HEP amplitude and RT in each subject. The thick yellow line indicates the mean slope of the linear regression averaged across subjects. (**a**) Correlations between HEP2 and RT at the occipitoparietal cluster for the High-fit group. (**b**) Correlations between HEP2 and RT at the occipitoparietal cluster for the Low-fit group.

## Discussion

We found significant differences in the neural processing of cardiac afferent signals in physically trained individuals, relative to their sedentary counterparts (Fig. 2a and 2b). Interestingly, our finding of lower HEP amplitude in the high-fit group at frontocentral sites is consistent with the result of decreased HEP at the same region after a session of heartbeat auditory feedback (Canales-Johnson et al., 2015). The pattern of HEP group differences in our study is very similar to the pattern reported by Canales-Johnson et al., with our high-fit group resembling their post-feedback condition (see Fig. 2a in comparison to Fig.2 B in Canales-Johnson et al. 2015). A lower HEP at frontocentral sites seems therefore to be indicating the effect of increased IS, acquired through learning. While Canales et al. induced learning using auditory feedback, our results suggest that regular physical exercise produces a similar effect.

The notion that physical exercise may increase IS is not new. Studies using cardiac perception have already shown that physically trained individuals demonstrate enhanced heartbeat awareness than those with less or no training (Jones and Hollandsworth, 1981; Schandry et al., 1993), while heartbeat awareness also improves during or immediately after a single session of physical exercise (Borg and Linderholm, 1967; Montgomery and Jones, 1984; Yuan et al., 2007; Rädler and Schandry, 1988). Our study, however, is the first to demonstrate this effect not using subjective reports of physical sensations, but through the measurement of a neurophysiological indicator bellow the threshold of consciousness. Thus, the methodology employed here captures primary aspects of interoception related to afferent signal transmission and its neural processing, and can help differentiate the concept of physiological IS from interoceptive awareness, which additionally requires a meta-cognition of internal body states (Schulz and Vögele, 2015).

Further, our results indicate that the increased IS observed in physically fit individuals during an attentional orienting phase induced by a warning signal, is associated with a quicker response to the posterior target stimulus (Fig. 3a). This effect is absent in the group of low-fit participants (Fig. 3b). This finding is consistent with previous research on interoception and cognition. Most studies in this area used subjective heartbeat detection tasks to prompt IS and found that increased interoception provides a cognitive advantage (Werner et al., 2010; Matthias et al., 2009; Dunn et al., 2010). Park et al., were the first to reveal a cognitive benefit associated to increased interoception indexed by the HEP. The authors showed that the HEP amplitude after a warning signal correlates with a successful visual discrimination of the upcoming stimulus (Park et al., 2014). Our study supports and extends this finding to a sustained attention task, designed to measure the capacity to effectively allocate attentional resources over time in order to react promptly to relevant stimuli. Sustained attention is ubiquitous in cognitive processing, since a reduced potential to monitor significant sources of information directly affects all cognitive abilities (Sarter et al., 2001), and is particularly critical for the performance of fundamental everyday activities (e.g. attending academic lessons at school: Suess et al. 1994, and driving: Larue et al. 2011), or highly responsible professional tasks (e.g. performing surgery: Di Stasi et al. 2014, piloting: Di Stasi et al. 2015, and handling air-traffic control: Loft et al. 2007). Linking the neural processing of visceral signals to such a crucial cognitive function is therefore elemental for the basic understanding of cognitive processing, and in particular, for elucidating the mechanisms that cause lapses of attention after focusing on a single task for an extended period of time.

Previous attempts to interpret the link between interoception and cognitive function pivot around the somatic marker hypothesis (Damasio et al., 1996), further postulating that the neural monitoring and representation of afferent body signals forms the basis for embodied self-awareness (Craig, 2002; Park et al., 2016; Babo-Rebelo et al., 2016). The sense of a physical “I” is a crucial requisite for goal-directed cognitive processing, and for successful behavioral and perceptual decision making. Our work puts forward the hypothesis that physical exercise leads to brain and cognitive improvements by enhancing embodied awareness. Earlier efforts to explain the fitness-brain-cognition relation focused on the effect of cardiovascular adaptations induced by physical exercise on CNS parameters, including cerebral blood flow and brain neurotrophic factors (e.g., BDNF and IGF-1; for a review see Hillman et al. 2008). Our results suggest that regular physical exercise additionally produces changes in the cortical processing of afferent cardiac information. This goes beyond the well-known structural and anatomical alterations in individual organ systems, and points towards complex brain-heart interactions as a mechanism contributing to enhanced cognition in physically fit individuals. Recent advances in the emerging field of network physiology promise therefore to find applications in cognitive neuroscience by offering a novel analytic and conceptual framework to investigate and understand how cognitive functions emerge out of multiple organ interactions (Bashan et al., 2012).

Further research is necessary to establish a robust link between physical fitness and neurophysiological markers of IS. Specifically, randomised control trials of fitness training programs should be performed to uncover a possible causal link, and to reveal if enhanced brain-heart communication linearly evolves as a result of increases in cardiovascular fitness, or whether other parameters, such as training type, intensity, duration, etc., come into play. Importantly, our findings open the way to study how other activities involving mind-body communication, for example dancing and interoceptive meditation, entrain brain-heart interactions with possible beneficial outcomes for cognitive function and performance.

On a methodological note, our study makes two significant contributions to the research on the HEP. First, our choice to use a cluster-based permutation test without an a priori definition of spatial or temporal regions of interest, made possible the discovery of a relevant occipitoparietal cluster that is being reported for the first time (for a review on HEP localisation in previous studies see Kern et al. 2013). Critically, the correlation between HEP amplitude and behavioral performance was found in this occipitoparietal cluster and not in the more popular frontocentral cluster (Fig. 2a vs. 2b). Second, group differences were only significant in the second heartbeat following the warning signal. This result highlights the importance of selecting an appropriate cardiac event to compute the HEP in future studies. An explanation of why we did not find significant group differences in the first post-warning HEP may relate to the short time interval between the occurrence of the first R-peak and the warning stimulus. Given the rapid, but not instantaneous, transmission of parasympathetic nerve traffic to the heart (Luque-Casado et al., 2016), it is likely that group differences in the neural response associated to the vagal output are not reflected in the first cardiac cycle following an attentional stimulus.

To summarise, the data we report here suggest that the regular practice of physical exercise changes how the brain processes transient afferent signals originating at the cardiovascular system. Further, our findings provide evidence that this differential neural processing is associated to enhanced cognitive performance, at least in high-fit individuals. These results offer neurophysiological support to the hypothesis that physical exercise improves cognitive function by modulating IS. Should this hypothesis is confirmed it will open the way to the design and development of novel cognitive training programs targeting directly the improvement of physiological interoception markers.

